# Distinct phenotypes and repertoires of bronchoalveolar and airway mucosal T cells in health and allergic asthma

**DOI:** 10.1101/2025.05.08.652962

**Authors:** Rod A. Rahimi, Neal P. Smith, Amandine Selle, Roya Best, Sidney Martin, Elizabeth Tuttle, Morris F. Ling, Benjamin D. Medoff, Alexanda-Chloé Villani, Andrew D. Luster

## Abstract

**Background:** T cells play a central role in host protection against respiratory pathogens, but a maladaptive T cell response can lead to pulmonary diseases. Defining the biology of protective versus pathogenic T cell responses in the lungs of humans will be critical to nominate novel approaches to improve respiratory health. Previous studies have examined T cells from the lungs captured via bronchoalveolar lavage (BAL), endobronchial brushings, or biopsies. However, whether these different approaches are capturing distinct T cell phenotypes and/or clonotypes remains unclear.

**Objective:** To evaluate and compare the transcriptional signatures of T cells isolated via BAL versus endobronchial brushings in healthy controls (HCs) and allergic asthmatics (AAs).

**Methods:** Flexible bronchoscopy was performed to obtain BAL and endobronchial brushings from 3 HC and 3 AA subjects. CD3+ T cells were sorted and single cell RNA- and T cell receptor (TCR)-sequencing was performed using the 10X Chromium platform. Unbiased clustering, differential gene expression analysis and TCR repertoire analysis was performed.

**Results:** Unbiased clustering analysis allowed us to define 7 CD8 and 6 CD4 T cell subsets. The most significant difference in T cell subsets abundance between AAs and HCs was the enrichment of CD4 T helper type 2 (Th2) cells when comparing endobronchial brush samples (OR=26.2, P=0.002), but not when examining BAL (OR=1.7, P=0.46), indicating differences in the T cell subsets captured from the BAL versus airway mucosa specimen processing. In further support of this observation, comparing the BAL and brush T cells across all subjects revealed an up-regulation of resident-memory T cell markers (i.e. *ITGAE, CD69*) in brush T cells versus BAL T cells in both CD4 and CD8 lineages. In contrast, BAL CD8 and CD4 T cells exhibited an enriched type I and II interferon signatures compared to brush T cells. Lastly, TCR repertoire analysis revealed that brush T cells contained dramatically expanded TCR clones. Expanded T cell clones from the brush expressed high levels of resident-memory markers, suggesting the airway mucosa is enriched for T_RM_ cells with unique TCR specificity.

**Conclusions:** Sampling T cells via BAL versus airway brushings yielded distinct T cell phenotypes and clonotypes with important implications for future research in lung immunology.

## Introduction

The lungs are exposed to a multitude of respiratory pathogens as well as noxious, non-infectious substances such as allergens (1,2). T cells play a central role in host protection from respiratory pathogens, but a maladaptive T cell response can lead to immunopathology (3,4). For example, in the context of respiratory viral infection, persistent T cell activation can drive tissue injury and fibrosis in murine models and is associated with severe viral pneumonia in humans (5,6). In addition, inappropriate CD4^+^ T helper type 2 (Th2) cell responses to aeroallergens orchestrate allergic inflammation in asthma (7). Defining the biology of protective versus pathogenic T cell responses in the lungs of humans will be critical to nominate novel approaches to improve respiratory health.

A variety of αβ T cells and unconventional T cell populations, such as gd T cells and MAIT cells, reside within the human lungs (8). Among the population of αβ T cells, there are different subsets of memory T cells, including tissue-resident memory T ( T_RM_) cells that are poised for recall responses, as well as distinct CD4^+^ T helper subsets (8,9). Previous studies have used bronchoalveolar lavage (BAL), endobronchial brushings, or biopsies to isolate T cells (10–16). However, whether these distinct approaches preferentially isolate different T cell populations and/or clonotypes remains unclear. For example, previous studies have yielded conflicting results regarding whether BAL and airway mucosal T cells exhibit similar or different expression of T_RM_ cell markers by flow cytometry, which may be due to tissue digestion altering cell surface proteins (15–17). Defining potentially distinct cellular features of BAL and airway mucosal T cells is necessary to define the optimal approaches to study lung T cells in different contexts.

Here, in healthy controls (HCs; n=3) and subjects with allergic asthma (AAs; n=3), we performed bronchoscopy with paired BAL and endobronchial brushings followed by cell sorting of CD3^+^ T cells and single-cell (sc) RNA- and TCR-sequencing analyses. The most significant difference in T cell subsets between AAs and HCs was the enrichment of T_H_2 cells when comparing endobronchial brush samples, but not when examining BAL, suggesting that T cells isolated from the BAL and airway mucosa exhibit distinct features. In further support of this concept, comparing the BAL and brush T cells across all subjects revealed an up-regulation of T_RM_ cell markers (i.e. *ITGAE, CD69*) in brush T cells whereas BAL T cells exhibited increased expression of genes involved in type I and II interferon signaling. Lastly, TCR repertoire analysis identified expanded T cell clones from the airway mucosa enriched in the expression of T_RM_ cell markers. In sum, these results suggest that T cell phenotypes and clonotypes are distinct in BAL and airway mucosal compartments. Our findings underscore the differences in sampling T cells from the BAL versus endobronchial brushings, providing an important technical resource for studying lung T cells in humans in the context of health and pulmonary diseases.

## Methods

### Subject recruitment

This study was a nonrandomized, nonblinded mechanistic study with the primary aim of comparing the transcriptional profiles of T cells from the airways of healthy controls (HCs) and allergic asthmatics (AAs). Volunteers were screened for eligibility with a full medical history, baseline spirometry, methacholine challenge testing, as well as allergen skin prick testing. In terms of inclusion criteria, male and female participants between 18 and 55 years old with either 1) no past medical history, normal spirometry and negative methacholine challenge test, and negative allergy skin prick testing (HCs) or (2) mild-to-moderate asthma with allergy, baseline FEV1 ≥ 75% of the predicted value and positive methacholine challenge, and positive skin prick test to cats or dust mites (AAs) were enrolled. A positive methacholine challenge test result was defined as a provocative concentration of less than or equal to 25 mg/ml causing a 20% drop in FEV1. AAs had positive skin prick testing to standardized HDM [*Dermatophagoides pteronyssinus*; Greer Laboratories, 10,000 allergy units (AU) per ml] and/or cat hair extract [Felis catis; Greer Laboratories, 10,000 bioequivalent allergy units (BAU) per ml]. Detailed inclusion and exclusion criteria can be found in Supplementary Materials, as previously published (12,14).

### Bronchoalveolar lavage and airway brushings

BAL samples were obtained by administering 120 ml of normal saline via the working channel of the bronchoscope into the right middle lobe (RML) with suctioning to collect the sample. After collecting the BAL, a 4-mm sterile nylon cytology brush was used to perform endobronchial brushing of a RML airway segment under direct visualization with a second cytology brush used to collect a second sample adjacent to the first endobronchial sample. The cytology brushes were each placed in RPMI media containing 5% human AB serum on ice prior to sample processing.

### Florescence-activated cell sorting and scRNA-seq

Cells were isolated from BAL and endobronchial brushings and stained with Fc receptor blockade, fluorescently-conjugated antibodies against CD45, CD14, CD20, CD3, and eFluor 780 fixable viability dye (Thermo Fisher Scientific). Additional information is provided regarding the fluorescently-conjugated antibodies used for florescence-activated cell sorting in table S1. A BD FACSAria Fusion flow cytometer (BD Biosciences) was used to sort eFluor 780-CD45+CD14-CD20-CD3+ cells. Analysis was performed with FlowJo v10.8 (Tree Star). Sorted cells were resuspended at a target concentration of 1000 cells per μl in preparation for loading on the 10X Chromium instrument (10X Genomics).

### Single-cell RNA and TCR libraries and sequencing

Single cell suspensions were profiled by scRNAseq with the Chromium Single Cell 5’ kit (V2, 10x Genomics PN-1000190). TCR-enriched cDNA libraries were generated with the Chromium Single Cell V(D)J Enrichment kit (10x Genomics PN-1000005). Library quality was assessed with an Agilent 2100 Bioanalyzer. All libraries were sequenced on an Illumina Nextseq 500/550 instrument. For the gene expression libraries, we used high output v2.5 75 cycles kits with the following sequencing parameters: read 1=26; read 2=46; index 1=10; index 2=10. For TCR libraries, we used 150 cycles kit with the following sequencing parameters : read 1=26; read 2=90; index 1=10; index 2=10.

### Read alignment and quantification

Raw sequencing data was pre-processed with CellRanger (v6.0.2, 10X Genomics) to demultiplex FASTQ reads, align reads to the human reference genome (GRCh38, v3.0.0 from 10X Genomics), and count unique molecular identifiers (UMI) to produce a *cell x gene* count matrix (30). All count matrices were then aggregated with Pegasus (v1.8.1, Python) using the *aggregate_matrices* function (31). Cells with >20% mitochondrial UMI or <500 unique genes detected were deemed low-quality cells and were filtered out of the matrix prior to proceeding with downstream analyses. The percent of mitochondrial UMI was computed using 13 mitochondrial genes (*MT-ND6, MT-CO2, MT-CYB, MT-ND2, MT-ND5, MT-CO1, MT-ND3, MT-ND4, MT-ND1, MT-ATP6, MT-CO3, MT-ND4L, MT-ATP8*) using the *qc_metrics* function in Pegasus. The counts for each remaining cell in the matrix were then log-normalized by computing the log1p(counts per 100,000), which we refer to in the text and figures as log(CPM).

### Cell clustering

For all cell clustering analysis, 2,000 highly variable genes were selected using the *highly_variable_features* function in Pegasus and used as input for principal component analysis. To account for technical variability between batches, the resulting principal component scores were aligned using the Harmony algorithm (32). The top 50 principal components were used as input for generating a neighborhood graph. Clustering the neighborhood graph was performed using the Leiden algorithm and the data was represented using the Uniform Manifold Approximation and Projection (UMAP) algorithm (spread=1, min-dist=0.5) (33,34).

### Marker gene identification

The marker genes defining each distinct cell cluster from our CD4 and CD8 T cell sub-clustering analyses were determined by applying two complementary methods. First, we captured genes with high expression in each cluster by calculating the area under the receiver operating characteristic (AUROC) curve for the log(CPM) values of each gene as a predictor of cluster membership using the *de_analysis* function in Pegasus. Genes with an AUC ≥ 0.75 were considered marker genes for a particular cluster. Second, we created a pseudobulk count matrix to identify genes with lower expression that were highly specific for a given cluster (35).

Specifically, we summed the UMI counts across cells for each unique cluster/sample combination to create a matrix of *n genes x (n samples*n clusters)* and performed “one-versus-all” (OVA) differential expression (DE) analyses for each cluster using the *DESeq2* package (v1.32.0, R v4.1.0) (36). For each cluster, we used an input model *gene ∼ in_clust*, where *in_clust* is a factor with two levels indicating if the sample was in or not in the cluster being tested. A Wald test was then used to calculate *P* values and compute a false discovery rate (FDR) using the Benjamini-Hochberg method. We identified marker genes that were significantly associated with a particular cluster as having an FDR<0.05. Top marker genes were visualized with the *ComplexHeatmap* package (v2.8.0, R) (37).

### Abundance analysis

To identify the association between cell subset abundance and either patient group (AA versus HC), we used a mixed-effects association logistic regression model similar to that described by Fonseka et al. (38). We used the *glmer* function from the lme4 package (v1.1-31, R) to fit a logistic regression model for each cell subset. Each subset was modeled independently with a “full” model as follows:

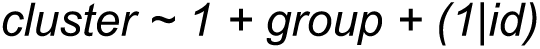

where *cluster* is a binary indicator set to 1 when a cell belongs to the given cell subset or 0 otherwise, *group* is a factor with 2 levels which represented the patient groups (AA or HC), and *id* is a factor indicating the donor. The notation *(1|id)* indicates that *id* is a random intercept. To determine significant associations, a “null” model of *cluster ∼ 1 + (1|id)* was fit and a likelihood ratio test was used to compare the full and null models. A false discovery rate was calculated using the Benjamini-Hochberg approach and clusters with an FDR<10% were considered significant.

### Differential gene expression analysis

Comparisons between groups and sample types were performed on pseudobulk count matrices using the *DESeq2* package (v1.32.0, R v4.1.0). The input model was either *gene ∼ group* (AA or HC) or *gene ∼ sample type* (brush or BAL). Significant DEG were identified using a Wald test (FDR<0.1).

### Gene set enrichment analysis

GSEA was performed using the *fgsea* function from the fgsea package (v1.18.0, R) with 10,000 permutations to test for independence. For GSEA performed to find gene sets associated with each sample type (brush, BAL), the input gene rankings were based on the pseudobulk log2FC values, where the gene with the highest value (brush-associated) was ranked first and the gene with the lowest value (BAL-associated) was ranked last.

### TCR analyses

A unique TCR clone was defined by the concatenation of the predicted V genes and CDR3 amino acid sequences from both the α and β chains. Diversity curves that measured Hill’s diversity metric across diversity orders 0–4 were created using the package alakazam (v1.0.2, R) with the alphaDiversity function (22).

### Data and code availability

scRNA-seq count matrices and related data will be deposited in the GEO database and raw human sequencing data will be available in the controlled access repository dbGaP upon publication of this study. Source code for all data analyses will be available on github upon publication of this study.

## Results and Discussion

### Tissue-resident T_H_2 cells are highly enriched in the airway mucosa, but not in the BAL, of subjects with allergic asthma

To define the similarities and differences in BAL versus airway mucosal T cells, we enrolled 3 healthy controls (HCs) adults and 3 adults with allergic asthma (AAs) (**Supplemental table 1**). For AAs, we enrolled individuals with a clinical diagnosis of mild asthma and defined allergy to house dust mites or cat, which was confirmed by skin prick testing. AAs had a FEV1 ≥ 75% predicted and positive methacholine challenge test, confirming the presence of airway hyper-responsiveness. Each subject underwent a flexible bronchoscopy and BAL. To sample T cells from the airway mucosa, we used two sequential endobronchial brushes to sample adjacent areas of third- to fourth-generation airway segments within the right middle lobe. We elected to sample the airway mucosa with brushes rather than endobronchial biopsies given the latter requires enzymatic tissue digestion, which has been shown to cleave T cell surface proteins and alter transcriptional signatures (17). In contrast, cells can be easily isolated from endobronchial brushes without the need for enzymatic tissue digestion (14). We sorted live CD45^+^CD14^-^CD20^-^ CD3^+^ T cells from the BAL and brushes, followed by paired scRNAseq and TCR-sequencing. using the 10X Chromium platform (**Fig. 1A**). We generated high-quality scRNA-seq data from 11 samples (5 BAL and 6 brush, totaling 78,602 cells) collected from three healthy controls (41,933 cells) and three allergic asthmatics (36,669 cells) (**Supplemental Fig. 1A; supplemental table 2**). Upon data integration and unsupervised clustering analysis (Materials and Methods), we identified 13 subsets grouped into major T cell lineages based on expression of the T cell co-receptors (*CD4, CD8A,* and *CD8B*) **(Fig. 1B).** When considering the relative abundance of the CD4 and CD8 lineages, we observed the BAL samples were enriched for CD4 T cells (log2FC = 0.46, p = 3.8e-5), while the brushings were enriched for CD8 T cells (log2FC = −0.27, p = 0.006) **(Supplemental Fig. 1B, supplemental table 3)**. We further sub-clustered 51,657 CD8 T cells and 26,897 CD4 T cells, generating 7 CD8 and 6 CD4 subsets respectively, which we annotated by cross-referencing top marker genes with published known T cell transcriptional signatures (AUC ≥ 0.75 and pseudobulk DGE FDR ≤ 0.05; Materials and Methods) (**Fig. 1C-D; Supplemental table 3)**. Among the CD8 subsets, we identified two T_RM_ subsets with cytotoxic features (CD8T*^GZMB,ITGAE^*; CD8T*^GZMK,EOMES^*), a lymph-homing subset (CD8T*^CCR7,SELL^*), two innate-like subsets (MAIT*^TRAV1-2, KLRB1^*; GDT*^TRDC,KIR2DL4^*), a cycling subset (CD8T*^STMN1, MKI67^*) and a high-mitochondrial subset (CD8T^high^ ^mito^). Among the CD4 subsets, we identified a T_H_1 subset (CD4T*^IFNG, CXCR6^*), a T_H_17-like subset (CD4T*^IL2, CCR6^*), a T_H_2 subset (CD4T*^IL13, IL17RB^*), a regulatory T cells (Tregs) (CD4T*^FOXP3, IL2RA^*), a lymph-homing subset (CD4T*^KLF2, SELL^*) and a subset defined by low genes (CD4T^low^ ^genes^). Whether the low-gene signature represents a quiescent T cell state or a technical artifact of cell quality remains unknown and requires additional future investigation. Among the CD4 T cells was also a small population of doublets, defined by the co-expression of the mast cell marker *CPA3* and the myeloid marker *LYZ*. First, we compared the relative abundance of each T cell subcluster between the BAL and brushes of all subjects combined. The CD8 T cell subsets enriched in the BAL included CD8T*^GZMK,EOMES^*(log2FC = 0.45, p = 0.001), CD8T*^CCR7,SELL^* ( log2FC : 0.49, p = 0.002), CD8T*^STMN1,MKI67^* (CD8 cluster 7 : log2FC = 0.75, p = 0.001) and MAIT*^TRAV1-2,KLRB1^* cells(log2FC = 0.4, p = 0.049) **(Fig. 1C, Supplemental table 4).** Among CD4 T cell populations, Tregs (log2FC = 0.32, p = 0.01) were enriched in the BAL **(Fig. 1D).** In contrast, CD4T*^IL2,CCR6^* were enriched in the endobronchial brushings (log2FC = −0.2, p = 0.001).

**Figure 1.**
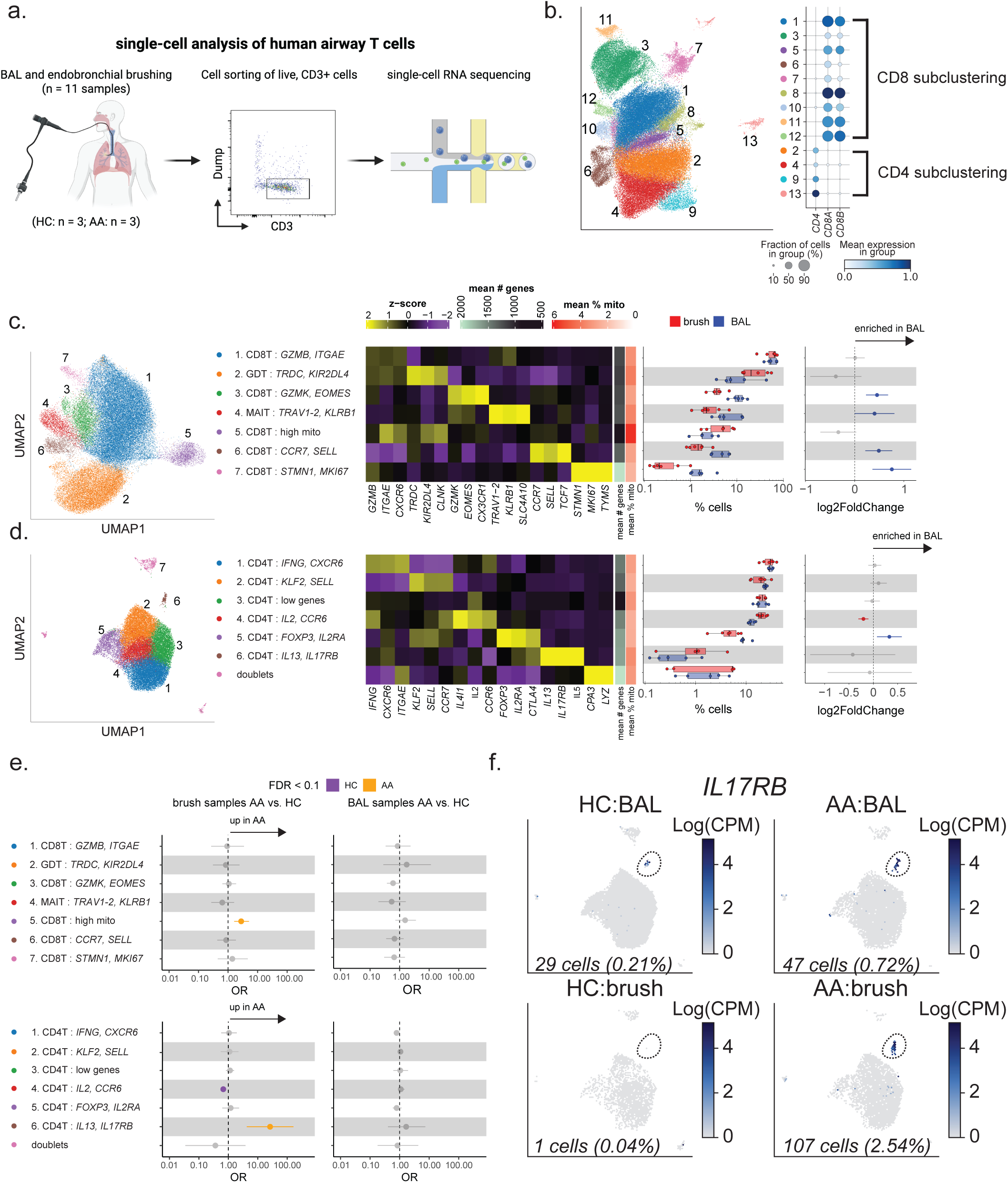
Tissue-resident Th2 cells are highly enriched in the airway mucosa, but not in the BAL, of subjects with allergic asthma. **(a)** Schematic of experimental design. **(b)** (left) UMAP embedding of 78,602 high-quality single cells, colored by predicted Leiden clusters. (right) Dot plot showing the percentage (size of the dot) and scaled expression (color) of *CD4, CD8A* and *CD8B* across the predicted clusters. Brackets indicate how clusters were segregated into CD4 and CD8 T cell sub-clusterings. **(c, d)** (left) UMAP embedding of CD4 and CD8 sub-clusterings, colored by predicted Leiden labels. (middle-left) Heatmaps showing select discriminative markers for each cluster when compared with every other cluster. Color scale denotes the normalized gene expression (mean zero, unit variance) for each cluster as well as the mean number of genes and percent mitochondrial UMIs. (middle-right). Abundance analysis comparing cellular frequencies of cell populations between brush (n = 6) and BAL (n = 5). Boxplots show the median (line) and IQR, with whiskers extending no more and 1.5X the IQR. (right) Dots represent log2FC between the brush and BAL population frequencies. Error bars represent 95% confidence intervals. Colored bars represent FDR < 0.1 (t-test). **(e)** Abundance analysis comparing cell frequencies between AA (n = 3) and HC (n = 3) in BAL and brush samples. Dots represent logistic regression odds-ratios. Error bars represent 95% confidence intervals. Colored bars represent FDR < 0.1. **(f)** UMAP Feature plots using color to indicate gene expression (logCPM) levels of *IL17RB* by HC (left) and AA (right) in BAL (top) and brush (bottom). Cell numbers and percentages represent gene expression across all CD4 T cells in the given group/sample combination.

Next, we sought to understand if there were differences in population abundance when comparing AA to HC in either the brush or BAL. When comparing the BAL samples, there were no significant differences between AA and HC **(Fig 1E, Supplemental table 5).** However, when comparing the endobronchial brush samples, we found 3 differentially abundant clusters. CD4T*^IL2,CCR6^* was enriched in HC (OR = 0.67, p = 0.003). CD8T^high^ ^mito^ was enriched in AA (OR = 2.86, p = 0.01). The greatest difference was the enrichment of the T_H_2 population (CD4T*^IL13,IL17RB^*) in the AAs exclusively in the brushings (OR = 26.2, p = 0.002), while that same effect was not seen in BAL (OR = 1.7, p = 0.46) **(Fig 1E-F).** These results indicate that the airway mucosa creates a unique cellular niche for maintaining T_H_2 cells in allergic asthma whereas the presence of T_H_2 cells within the BAL does not correlate with clinically significant airway disease. Although allergens are known to induce FOXP3^+^ Treg responses, we did not find differences in abundance of Tregs between HCs and AAs in either the BAL (OR = 0.8, p = 0.3) or brushes (OR = 1.2, p = 0.57) (**Fig. 1E**), suggesting the niche supporting T_H_2 cells may not maintain allergen-specific FOXP3^+^ Tregs to the same degree (18,19). Altogether, our results demonstrate that BAL and endobronchial brushings yield distinct T cell populations, including T_H_2 cells in the context of allergic asthma.

### Airway mucosal T cells exhibit an enriched tissue-resident memory signature whereas BAL T cells exhibit an enriched interferon signature

We next sought to comprehensively examine the expression of marker genes that are known to be upregulated in T_RM_ cells across our defined T cell subsets, including *ZNF683*, *RUNX3*, *PRDM1*, *PDCD1*, *JUNB*, *ITGAE*, *CXCR6*, *CD69*, and *CD101* **(Supplemental Fig. 1C-D, supplemental table 6)** (8). We observed broad expression of most of these T_RM_ markers among our CD8 populations, with the highest expression in CD8T*^GZMB,ITGAE^*, GDT*^TRDC,KIR2DL4^*, CD8T^high^ ^mito^ and CD8T*^STMN1,MKI67^.* Overall, the expression levels of these genes were lower but present in our CD4 populations, with the T_H_1 population (CD4T*^IFNG,CXCR6^*) having the highest expression levels of the integrins *ITGAE* and *ITGB1.* We next compared the expression of these T_RM_-related genes in our BAL versus brush T cells across the defined CD4 and CD8 subsets. Compared to BAL T cells, the brush CD8 and CD4 T cell clusters exhibited a greater expression in T_RM_ cell markers with these genes being most upregulated in *GZMB*-expressing CD8 T cells and Th1 cells from the brush (**Fig 2A-B**). Expression of *ITGAE* and *CD69*, which represent the most canonical T_RM_ cell markers, were higher in brush CD8 and CD4 T cells in both HCs and AAs, underscoring that airway brushings yield T cells with an enriched T_RM_ cell signature compared to BAL T cells regardless of disease status (**Fig 2B**).

**Figure 2.**
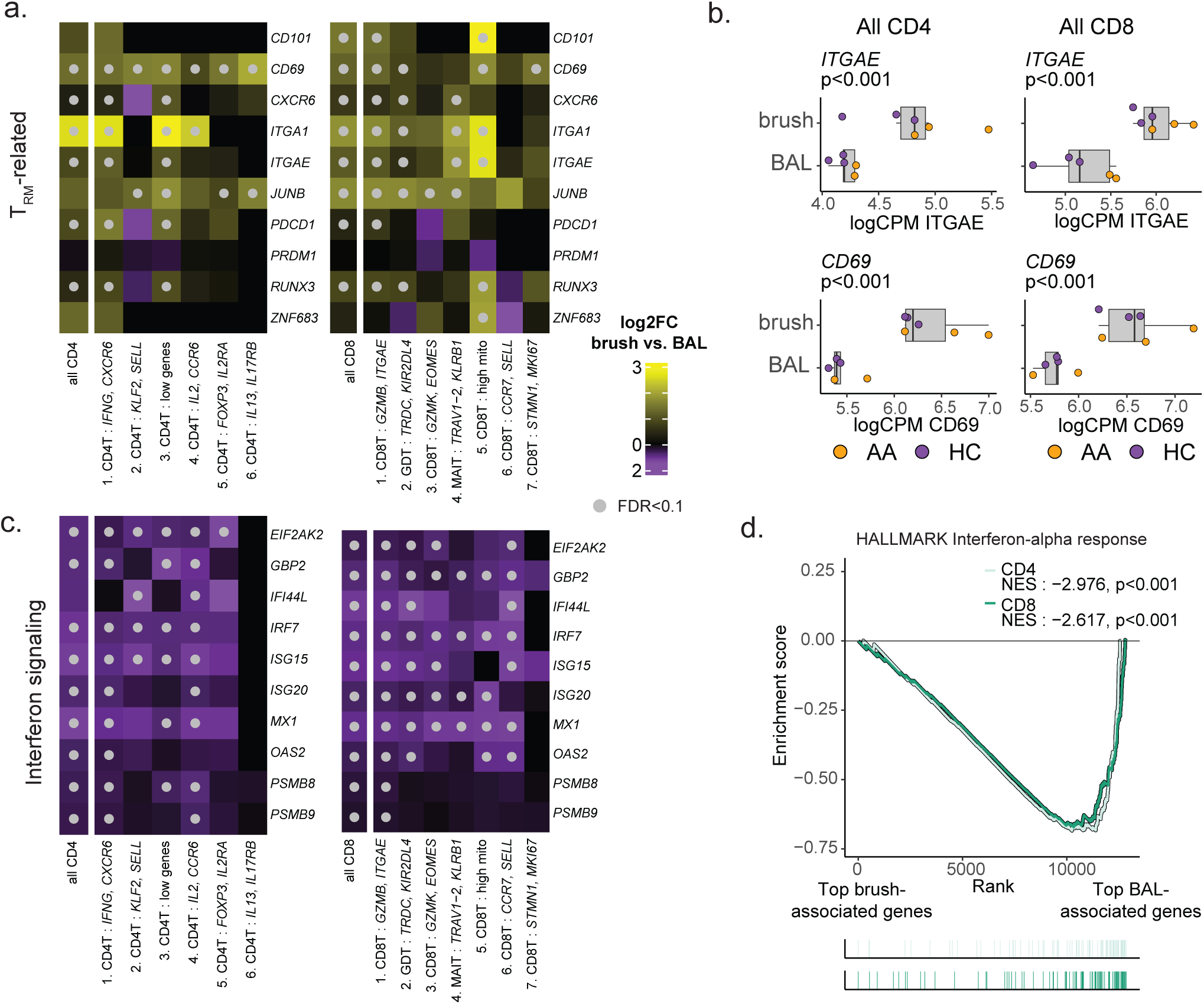
Airway mucosal T cells exhibit an enriched T_RM_ cell signature whereas BAL T cells exhibit an enriched interferon signature. **(a, c)** Heatmaps showing the differential gene expression results (brush versus BAL) for **(a)** known T_RM_ genes and **(c)** select interferon-stimulated genes across each (left) CD4 and (right) CD8 cluster as well as each lineage in aggregate (“all CD4”, “all CD8”). Color scale indicates the log2FC difference between brush and BAL samples. Grey dots represent FDR < 0.1 (Wald test). **(b)** logCPM of *ITGAE* and *CD69* for cells in the (left) CD4 and (right) CD8 lineages between brush and BAL samples. Color indicates the patient group the sample belonged to. **(d)** GSEA of the HALLMARK interferon alpha response gene signature for both the CD4 and CD8 lineages.

Following our analysis of T_RM_ cell markers, we focused on identifying pathways more highly expressed in BAL versus brush T cells. We noted markedly higher expression of interferon (IFN) genes in BAL versus brush T cells across most populations, with the notable exception of T_H_2 cells (CD4T*^IL13,IL17RB^*) (**Fig 2C**). We performed gene set enrichment analysis (GSEA), which identified type I interferon responses being strongly associated with both CD4 and CD8 BAL cells **(Fig 2D, Supplemental Fig. 1E-F, supplemental table 7).** In sum, our findings underscore the phenotypic differences between BAL and brush T cells in humans.

### T cell clones are enriched in the airway mucosa

Given our demonstration of phenotypic differences between BAL and brush T cells, we aimed to determine whether the clonotypes in these two compartments were distinct by analyzing the TCR repertoires of our captured T cells. scTCRseq allowed us to capture fully resolved TCRs from approximately 55.3% of the T cells in our data, with a capture rate fairly uniformed across samples **(Supplemental Fig. 1G)**. Using Hill’s diversity index, which captures both the richness (low q values) and abundance (high q value) of the TCR repertoire, we found that the BAL samples were significantly more diverse than brush samples (**Fig 3A**) (22). We subsequently compared the relative proportions that the expanded TCR clones (> 0.5% of a donor’s captured repertoire) represented in the BAL and brush **(Fig 3B).** There were 30 TCRs with >2 fold-increase in the brush repertoire when compared to the BAL, of which 27 were found to be CD8 T clones (**Fig 3B**). Interestingly, we found strong correlations between the clones’ relative abundance in brush versus BAL and the expression of T_RM_ markers *ITGAE* (r = 0.54, p<0.001)*, CD69* (r = 0.63, p<0.001)*, ITGA1* (r = 0.75, p<0.001) and *CXCR6* (r = 0.35, p = 0.009) **(Fig 3C).** This suggests the airway mucosa is enriched for T_RM_ cells with unique TCR specificity, presumably recognizing previously encountered respiratory pathogens.

**Figure 3.**
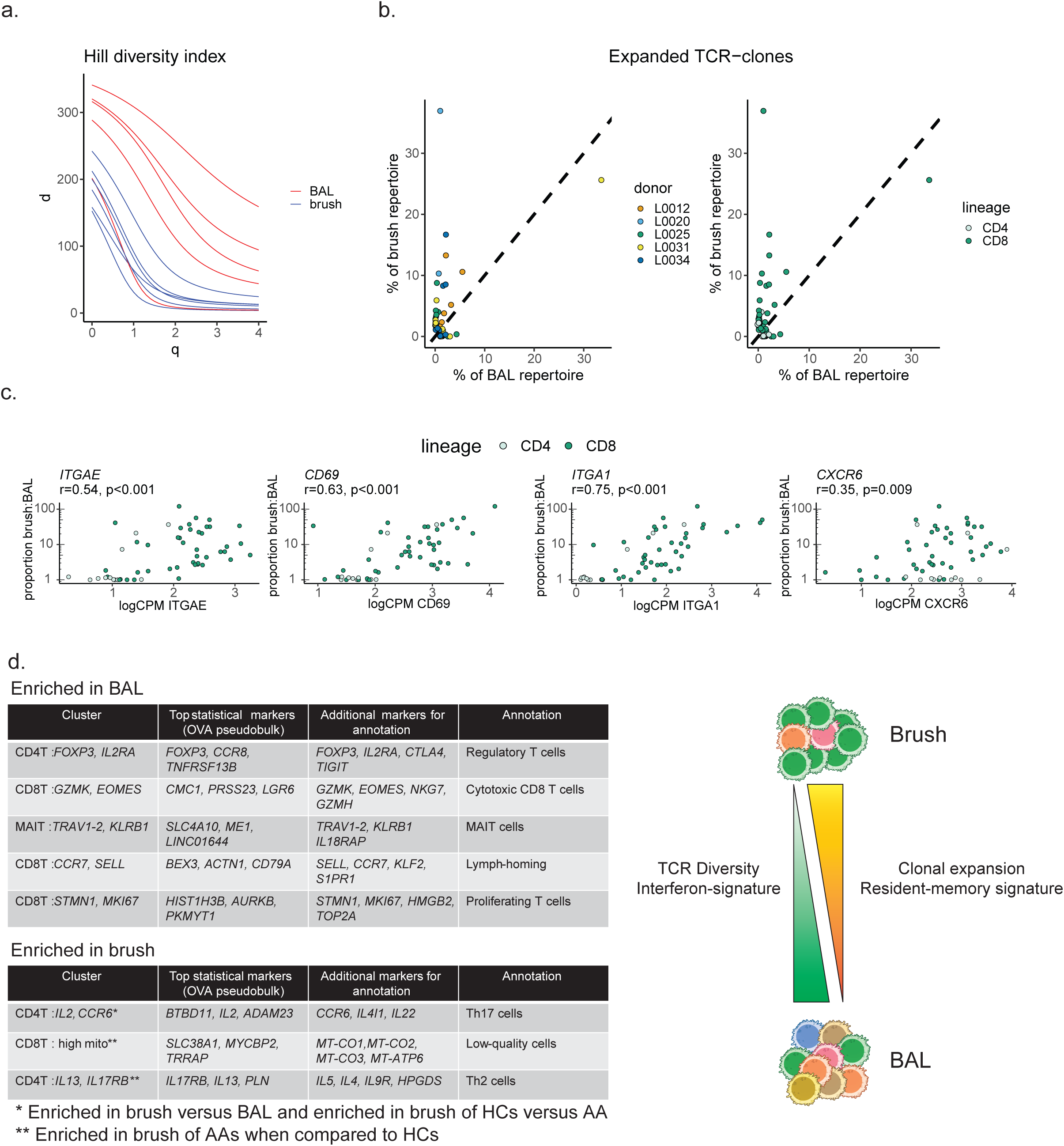
T cell receptor clones enriched in the airway mucosa. **(a)** Smoothed Hill’s diversity curve at diversity orders 0–4 for BAL (red) and brush (blue) samples. **(b)** Each dot corresponds to an expanded clone with the x- and y-axes showing the proportion of the BAL and brush repertoires that each clone represents. Each clone is colored by donor (left) and the lineage it came from (right). **(c)** Each dot corresponds to an expanded clone, with the y-axis representing the clone’s proportion in the brush relative to BAL and the x-axis representing the mean logCPM expression of T_RM_ genes *ITGAE, CD69, ITGA1, CXCR6.* Statistics represent those from Pearson correlations. **(d)** Summary of major findings. (left) table listing cell subsets that were enriched either in the BAL (top) or brush (bottom). Included are the genes used for annotating the clusters and our predicted annotation. (right) Summary of major phenotypic and TCR repertoire differences in brush and BAL.

Our study comparing the transcriptional and clonal signatures of BAL and endobronchial brush T cells in healthy controls and allergic asthmatics yields several important findings. First, although BAL is often used to sample immune cells from the lungs, we show that the endobronchial brush samples, but not the BAL, were enriched for T_H_2 cells in allergic asthma. Such an observation may explain the previous finding that BAL-derived immune cells failed to identify an enrichment of T_H_2 cells in allergic asthma compared to controls (11). Second, T cells sampled by airway brushing exhibited up-regulated T_RM_ cell markers whereas BAL T cells demonstrated a more pronounced interferon signature. Although the mechanism and role of the IFN signature in BAL T cells will require additional study, given that alveolar macrophages are a dominant producer of type 1 interferons and a subpopulation of alveolar macrophages exhibit a basal IFN signature in healthy individuals, it is possible that basal interferon production from alveolar macrophages maintains BAL T cells in a state poised for anti-viral responses (20,21). Third, the TCR repertoires of BAL and brush T cells are distinct, with higher diversity among BAL T cells and certain clonotypes being dramatically enriched in brush samples. While defining the specificity of these expanded T cell clones from brush samples will require future study, it underscores that unique T cells are recovered from endobronchial brushing compared to BAL. One limitation of our study is that the endobronchial brushes are sampling third- to fourth-generation airway segments, and we were unable to sample airway mucosal T cells within more distal airways. Future studies will be necessary to define the differences in airway mucosal T cells from the proximal to distal airways, potentially using deceased donor lungs. Our findings support data from murine models in which the airways provide niches to support T_RM_ cells with CD8^+^ T_RM_ cells being abundant within the airway epithelium and CD4^+^ T_RM_ cells being maintained in adventitial niches surrounding airways or within inducible bronchus-associated lymphoid structures (23–29). Lastly, our approach of using two endobronchial brushes followed by enrichment of CD3^+^ T cell via cell sorting provides a technical approach to obtain a sufficient number of T cells from the airway mucosa for downstream analysis while avoiding tissue digestion. In sum, our study characterizes important, distinguishing features of lung T cells isolated via BAL versus endobronchial brushing, providing a technical resource for studying T cells in the human lungs in health and different disease states.

## Acknowledgments

We thank the staff of the Massachusetts General Hospital Pulmonary Special Procedures Unit and Translational Clinical Research Center as well as the clinical research coordinators (M. Kone, G. Lopes).

---

**Supplemental figure 1.**
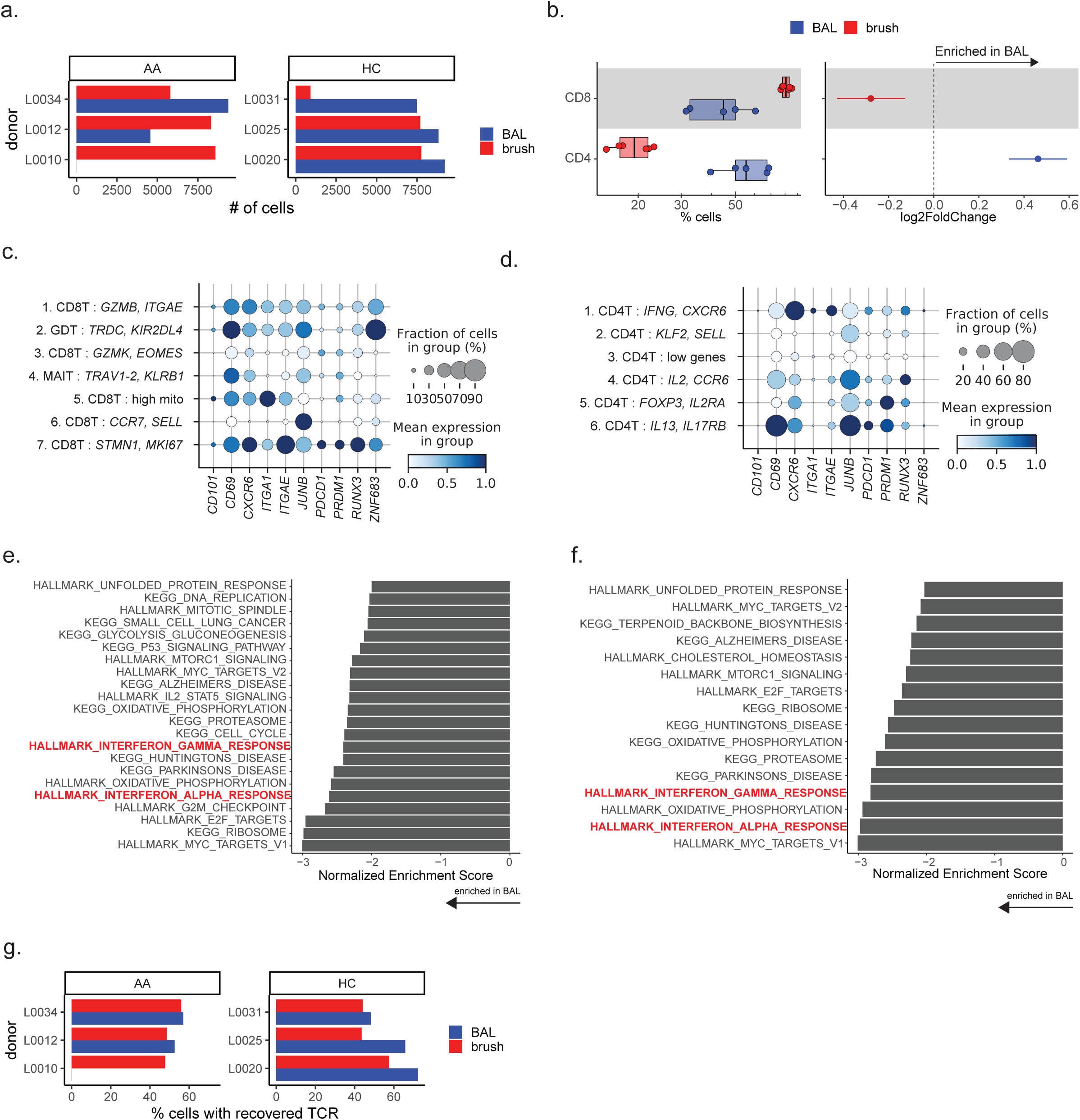
**(a)** The number of high-quality cells recovered from each sample. **(b)** abundance analysis comparing frequencies of CD4 and CD8 T cells between brush (n = 6) and BAL (n = 5). (left) boxplots show the median (line) and IQR, with whiskers extending no more and 1.5X the IQR. (right) Dots represent log2FC. Error bars represent 95% confidence intervals. Colored bars represent FDR < 0.1. **(c-d)** Dot plot showing the percentage (size of the dot) and scaled expression (color) of T_RM_ markers across the CD8 (left) and CD4 (right) clusters. **(e-f)** Gene set enrichment analysis (GSEA) of DE results comparing brush versus BAL cells from the CD8 (le) and CD4 (right) lineages. **(g)** The percentage of cells with a recovered TCR across samples.

**Supplemental Table 1. Study participant characteristics.** Clinical data for the participants in the study. Each row is a patient (ID) and information on which clinical group (Group), biological sex, age, race, ethnicity, allergen sensitization, forced expiratory volume in 1 second (FEV1), forced vital capacity (FVC) and which samples were collected (BAL sample, Brush sample) are included.

**Supplemental Table 2. scRNAseq QC and sample information.** Each row represents a sample (Channel) and included is information regarding the mean # of genes, mean percent mitochondrial UMIs, the number of cells that passed our QC threshold, the library primers used for gene expression libraries (gex_index) and TCR libraries (TCR_index).

**Supplemental Table 3. Marker genes for cell subsets.** Each row represents a marker gene (featurekey) for each cellular subset with corresponding AUROC values (AUROC), one-vs.-all (OVA) pseudobulk *P* value (OVA pseudobulk pval), OVA pseudobulk FDR (OVA pseudobulk padj), and OVA pseudobulk log2(fold change) (OVA pseudobulk log-fc), marker gene significance as determined by AUROC (AUROC_marker) and pseudobulk (pseudobulk_marker) approaches, cell subset (annotation) and the corresponding lineage (lineage). Genes with an AUC ≥ 0.75 or a pseudobulk FDR < 0.05 are included for each subset.

**Supplemental Table 4. BAL versus brush abundance statistics.** Each row is a cell subset (annotation) in a given lineage (lineage) and included are the T-test statistics for comparing proportions from BAL and brush samples. These statistics include the log2 fold-change (log2FoldChange), T-statistic(statistic), lower- and upper-limits of the 95% confidence interval (conf.low, conf.high), p-value (p.value), FDR-adjusted p-value (padj) and whether or not it met our statistical significance criteria (significance).

**Supplemental Table 5. AA versus HC abundance statistics.** Each row corresponds to a comparison between AA and HC from a specific sample type (sample_type) in a given cell subset (annotation) and lineage (lineage). Statistics included are the p-value (model.pvalue), odds-ratio (groupAA.OR), upper- and lower-limits of the 95% confidence interval (groupAA.OR.95pct.ci.lower, groupAA.OR.95pct.ci.upper) and FDR-adjusted p-value (padj).

**Supplemental Table 6. Differential gene expression.** Statistics for differential gene expression analysis comparing BAL versus brush samples. Information included is the gene name (gene), normalized transcript counts averaged for all samples (baseMean), log2 fold-change (log2FoldChange), standard error (lfcSE), Wald-statistic (stat), P value (pvalue), FDR-adjusted p-value (padj), the cell subset for the comparison (cluster) and the corresponding lineage (lin).

**Supplemental Table 7. Gene set enrichment analysis comparing brush and BAL.** Statistics for GSEA on the DEGs comparing brush to BAL samples. Each row is a gene set tested (pathway) in a given subset (cluster) from a specific lineage (lin) and included are the associated p-value (pval), FDR (padj), enrichment score (ES), normalized enrichment score (NES), the number of times a random gene set had a more extreme enrichment score value (nMoreExtreme), number of genes from the gene set found in the cellular subset (size), and the leading edge (leadingEdge).

